# Atypical speech production of multisyllabic words and phrases by children with developmental dyslexia

**DOI:** 10.1101/2022.08.24.505144

**Authors:** Mahmoud Keshavarzi, Giovanni M. Di Liberto, Fiona Gabrielczyk, Angela Wilson, Annabel Macfarlane, Usha Goswami

## Abstract

The prevalent ‘core phonological deficit’ model of dyslexia proposes that the reading and spelling difficulties characterizing affected children stem from prior developmental difficulties in processing speech sound structure, for example perceiving and identifying syllable stress patterns, syllables, rhymes and phonemes. Yet spoken word production appears normal. This suggests an unexpected disconnect between speech input and speech output processes. Here we investigated the output side of this disconnect from a speech rhythm perspective by measuring the speech amplitude envelope (AE) of multisyllabic spoken phrases. The speech AE contains crucial information regarding stress patterns, speech rate, tonal contrasts and intonational information. We created a novel computerized speech copying task in which participants copied aloud familiar spoken targets like “Aladdin”. Seventy-five children with and without dyslexia were tested, some of whom were also receiving an oral intervention designed to enhance multi-syllabic processing. Similarity of the child’s productions to the target AE was computed using correlation and mutual information metrics. Similarity of pitch contour, another acoustic cue to speech rhythm, was used for control analyses. Children with dyslexia were significantly worse at producing the multi-syllabic targets as indexed by both similarity metrics for computing the AE. However, children with dyslexia were not different from control children in producing pitch contours. Accordingly, the spoken production of multisyllabic phrases by children with dyslexia is atypical regarding the AE. Children with dyslexia may not appear to listeners to exhibit speech production difficulties because their pitch contours are intact.

**Research Highlights:**

- Speech production of syllable stress patterns is atypical in children with dyslexia.
- Children with dyslexia are significantly worse at producing the amplitude envelope of multi-syllabic targets compared to both age-matched and reading-level-matched control children.
- No group differences were found for pitch contour production between children with dyslexia and age-matched control children.
- It may be difficult to detect speech output problems in dyslexia as pitch contours are relatively accurate.

## 1. Introduction

Developmental dyslexia is a disorder of learning that primarily affects accurate and fluent word reading and spelling (Lyon, Shaywitz, & Shaywitz, 2003). Although there are multiple theories regarding etiology, including theories based on atypical visual processing (Stein and Walsh, 1997), atypical spatial attention (Facoetti et al., 2010) and atypical cerebellar function (Nicolson, Fawcett & Dean, 2001), the only theory with supporting data from infancy onwards is the phonological theory (Goswami, 2022, for a recent theoretical review). The phonological ‘core deficit’ theory (Stanovich, 1988) proposes that the core problem for affected children involves achieving cognitive awareness of word sound structure. At the level of sensory/neural processing of speech, this core phonological deficit could arise either from atypical perception of the speech signal, atypical production of the speech signal, or (more likely given the developmental connection between perception and production) *both* atypical perception and production. The oral tasks testing phonological theory have typically been perception tasks. Children with dyslexia have difficulty in perceiving whether words rhyme, difficulty identifying the mis-stressing of multi-syllabic words (prosodic difficulties), difficulty counting syllables in words, and difficulty counting or manipulating phonemes (the smaller sound elements typically represented by the alphabet) (Ziegler & Goswami, 2005). These perceptual difficulties are found across spoken languages, whether oral syllable structure is simple (that is consonant-vowel, CV, as in Spanish) or complex (CCVCC, as in English) (Elbro, Borstrøm, & Petersen, 1998; Law, Wouters, & Ghesquiere, 2017; Ziegler et al., 2010). In the cross-language speech processing literature, impaired perception of phonology typically relates to impaired phonological production. For example, native speakers of Japanese have difficulty in perceiving the difference between the English speech sounds associated with the letters L and R, and also have difficulty in producing this phonological difference (Bradlow, Pisoni, Akahane-Yamada, & Tohkura, 1997). Yet despite their difficulties in perceiving the phonological structure of words, difficulties in *producing* accurate phonological structures have been difficult to demonstrate in children with dyslexia. This has led some researchers to propose that the phonological difficulties in dyslexia are primarily related to ‘input phonological representations’ – the representations required for speech perception rather than for speech production (Szenkovits & Ramus, 2005).

However, for very young children there is evidence for speech production difficulties in dyslexia. In one longitudinal sample (Smith, Smith, Locke, & Bennett, 2008), toddlers who were genetically at-risk for dyslexia via family risk showed greater pausing and reduced production of syllables per second at 2 years of age compared to not-at-risk toddlers, suggestive of difficulties in preparing multi-syllabic utterances. In earlier assessments between 8 – 19 months, the same at-risk toddlers had shown atypical babbling and proto-speech, producing less complex syllable structures, an early indicator of reduced phonological sophistication (Lambrecht Smith, Roberts, Locke, & Tozer, 2010). In a separate line of research, word finding difficulties in tasks requiring picture naming of familiar items (‘confrontation naming’ tasks) have frequently been documented in children with developmental dyslexia (Constable, Stackhouse, & Wells, 1997; Denckla & Rudel, 1976; Goswami, Schneider, & Scheurich, 1999; Katz, 1986; Snowling, Van Wagtendonk, & Stafford, 1988; Swan & Goswami, 1997; Wolf & Goodglass, 1986), particularly for multisyllabic items, whether of high frequency (e.g., *television*) or low frequency (e.g., *binoculars*). Producing the names for multi-syllabic items is significantly impaired when children with dyslexia are compared to reading-level-matched (RL) control children in addition to chronological-age-matched (CA) control children. The RL-matched children are younger than the children with dyslexia and consequently have less oral language learning and a lower mental age. When children with dyslexia perform more poorly than younger children despite being matched on word recognition and having a higher mental age and greater oral language experience, this is important in helping to establish potential causal factors (Bryant & Goswami, 1986). Speech production would be expected to become more adult-like with greater oral experience. Speech production would also be expected to become more adult-like with greater reading experience, as reading acquisition is known to fundamentally change speech processing for all learners (Ziegler & Ferrand, 1998). Accordingly, if speech production by children with dyslexia was less accurate even than that of younger RL-matched children, for example regarding a parameter like amplitude envelope (AE) similarity or pitch contour similarity, this would support the view that atypical sensory/neural processing of either the speech AE or of pitch-related parameters respectively might be a causal factor in dyslexia.

Dyslexic difficulties in accessing the aspects of the phonological structure of words that are dependent on the speech AE would be predicted on the basis of the Temporal Sampling (TS) theory of developmental dyslexia (Goswami, 2011), an auditory theory based on impaired perception of the speech AE from infancy onwards (see Goswami, 2022, for infant data). The speech AE is a power-weighted average of the amplitude modulations (or speech energy fluctuations) produced at different temporal rates by speakers of human languages (Varnet et al., 2017). TS theory was developed to explain a decade of research studies across languages that indicated impaired perception of AE rise times and impaired amplitude modulation detection for children with dyslexia (e.g., Lorenzi et al., 2000; Goswami et al., 2002; see Goswami, 2011, 2015 for reviews). The perceptual effects of rise times in the AE help to create speech rhythm, as when we deliberately speak to a rhythm, the rise times of the vowels in stressed syllables are produced approximately isochronously (Scott, 1998; Goswami & Leong, 2013). AE rise times are also important sensory cues to stress placement in multi-syllabic words, as larger rise times at lower frequencies in the AE help to indicate stressed syllables (Greenberg, 2006). More recently, work in adult auditory neuroscience has indicated AE rise times as a key property in the neural processing of speech. Cortical signals reliably encode acoustic AE information (Lalor et al., 2009; Daube et al., 2019) and, from a neural oscillations perspective, AE rise times trigger oscillatory phase resetting (Giraud & Poeppel, 2012). Adult neural data indicate that these oscillatory processes are a key aspect of speech encoding (Gross et al., 2013). The speech signal is encoded cortically by oscillatory neural networks which exhibit intrinsic rates similar to the rhythmic rates of different patterns of amplitude modulation nested in the speech signal. TS theory proposes that the automatic alignment of these endogenous brain rhythms with rhythm patterns in speech is atypical in children with developmental dyslexia from birth, in part because the sensory cues (AE rise times) that trigger automatic neural alignment are perceived poorly by infants at family risk for dyslexia (Kalashnikova et al., 2018), and are perceived poorly by children with dyslexia (e.g., Goswami et al., 2011). Children with dyslexia also show impaired neural representation of low frequency AE information (Power et al., 2016; Molinaro et al., 2016). As the AE is perceived and represented poorly by children with developmental dyslexia, it is logical to expect that the AE might also be *produced* poorly during natural speech.

To explore this hypothesis, we created a novel syllable stress interface for children which enabled us to measure different features of their natural speech production. The task was based on copying adult-produced oral target words, which were spoken as names for pictures of familiar child words such as “Aladdin”. We then tested both children with dyslexia with this computerized interface and CA- and RL-matched controls. Based on TS theory, we expected the children with dyslexia to produce AEs for target items that were dissimilar to the target, while children without dyslexia should show speech production of AEs that were more similar to the target. For example, children with dyslexia might be expected to produce inconsistent differences in amplitude between strong and weak syllables, resulting in the overall envelope of their speech production being substantially different from the target. Children heard the computer speak a pictured known word (e.g. ‘Aladdin’) and simultaneously saw the target AE as a line on the computer screen (see Figure 1). They were asked to repeat the target word 3 times, simultaneously viewing their own AE, which appeared in real time overlaying the target AE on each occasion to provide visual feedback. Children were encouraged to try to match their AE to the target AE. The online interface automatically computed the averaged child AE, simultaneously computing how well the pitch contour matched the target pitch contour (pitch contour was not depicted on-screen). Pitch contour was selected as a control feature as it also captures the acoustic changes as speakers move from producing strong to weak syllables, and is known linguistically as an important aspect of speech processing (Lieberman, 1965). This ‘syllable stress game’ was administered to 75 children aged between 9 and 11 years who either met criteria for developmental dyslexia or who showed typical reading development.

**Fig 1.**
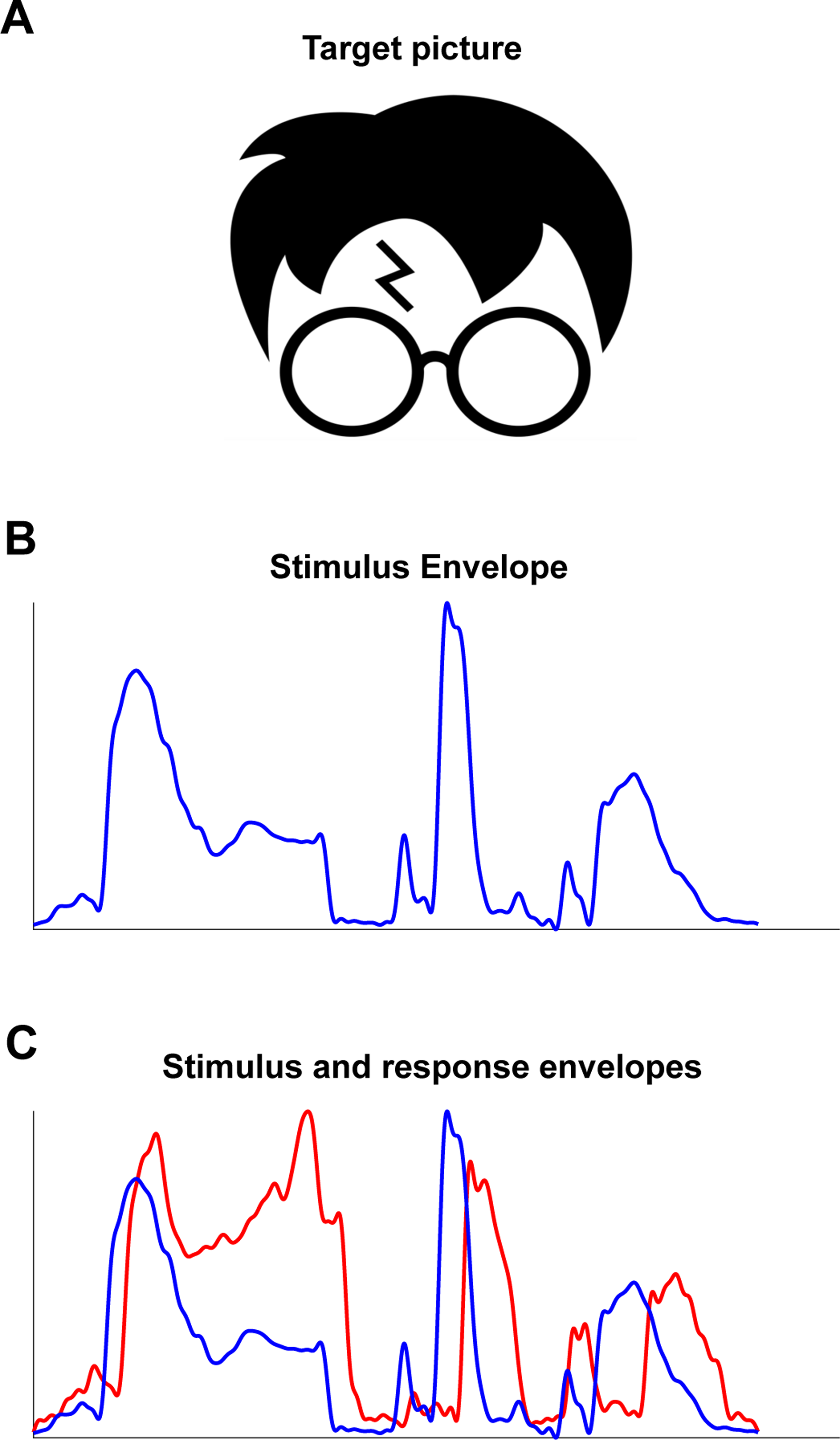
Schematic depiction of sequence of events during a single trial for the speech stimulus “Harry Potter”. The child first saw a picture related to the target stimulus (panel A), then a depiction of the stimulus envelope as the target word was pronounced (panel B, unfolding in real time). Although not depicted for the child, the horizontal axis in panels B and C is time and the vertical axis is amplitude. The stimulus envelope (blue) remained visible and the participant’s response envelope was overlaid as they finished speaking (panel C). The child was instructed to repeat the target stimulus in a time duration of 3 seconds. As the child repeated the stimulus, the envelope of the target stimulus with the child’s response envelope overlaying were shown on each occasion on the main layout (panel C).

## 2. Materials and methods

### 2.1. Participants

Seventy-five children aged between 9 and 11 years participated: 19 chronological-age-matched control children (CA, average age 11 years 0 months), 20 reading-level-matched control children (RL, average age 9 years 4 months), 19 children with dyslexia who were receiving an oral rhythmic intervention comprising 18 sessions of musical and rhythmic language activities spread over the school term (DY1, average age 11 years 4 months), and 17 children with dyslexia who were awaiting intervention (DY2, average age 10 years 8 months). The task was given as an add-on to the intervention for the DY1 children, but in contrast to the other oral rhythmic tasks being used in the intervention, the experimenter did not provide feedback nor help the child to perform the task. All participants were taking part in an ongoing longitudinal study of auditory processing in dyslexia (Keshavarzi et al., 2022a; Keshavarzi et al., 2022b; Mandke et al., 2022). Unfortunately the original study design (group matching based on data collected in 2017-18) was impacted by the Covid pandemic, which began closing UK schools in March 2020. At the current test point (Spring 2022), the remaining participants in the DY1 group were significantly older than the CA and DY2 groups, despite the groups being well-matched in age when the study began. Note that although this age difference could confound group comparisons (as the DY1 children have the advantage of being older than their age-matched controls), the age difference goes against our *a priori* hypothesis that the DY1 children will actually be worse than the CA children regarding AE production. Being older, the DY1 group will have more oral language experience than the CA children. Accordingly, if maturation was the sole driver of how adult-like the AE of children’s speech becomes, then the DY1 group should be at an advantage rather than a disadvantage compared to the CA children concerning the adult-likeness of their speech production.

All participants had English as their native language, and had previously received a short hearing test across the frequency range 0.25 – 8 kHz (0.25, 0.5, 1, 2, 4, 8 kHz) using an audiometer, and were sensitive to sounds within the 20 dB HL range. For DY1, the syllable stress production data were collected during sessions 7 and 9 of the 18-session oral rhythmic intervention. Participant characteristics by group are shown in Table 1. Oral language as measured by the British Picture Vocabulary Scales (Dunn & Dunn, 2009) did not differ between groups. Intelligence (IQ) was pro-rated from 4 subtests of the Wechsler Intelligence Scale for Children administered when the project began (WISC (Wechsler, 2016); similarities, vocabulary, block design and matrix reasoning) and did not differ between groups. Nonverbal IQ (Matrix subtest of the WISC, scaled score) reassessed closer to the time of receiving the speech production task also did not differ between groups, see Table 1. However, performance in reading (British Ability Scales [BAS (Elliot, Smith, & McCulloch, 1996)] single word reading and Test of Word Reading Efficiency [TOWRE (Torgesen, Wagner, & Rashotte, 1999)] nonword reading), BAS spelling and phonology (Phonological Awareness Battery [PhAB] rhyme and phoneme awareness (Frederickson, Frith, & Reason, 1997) was significantly lower for the DYS participants. All children and their parents gave informed consent for the study in accordance with the Declaration of Helsinki, and the study was approved by the Psychology Research Ethics Committee of the University of University.

**Table 1.**
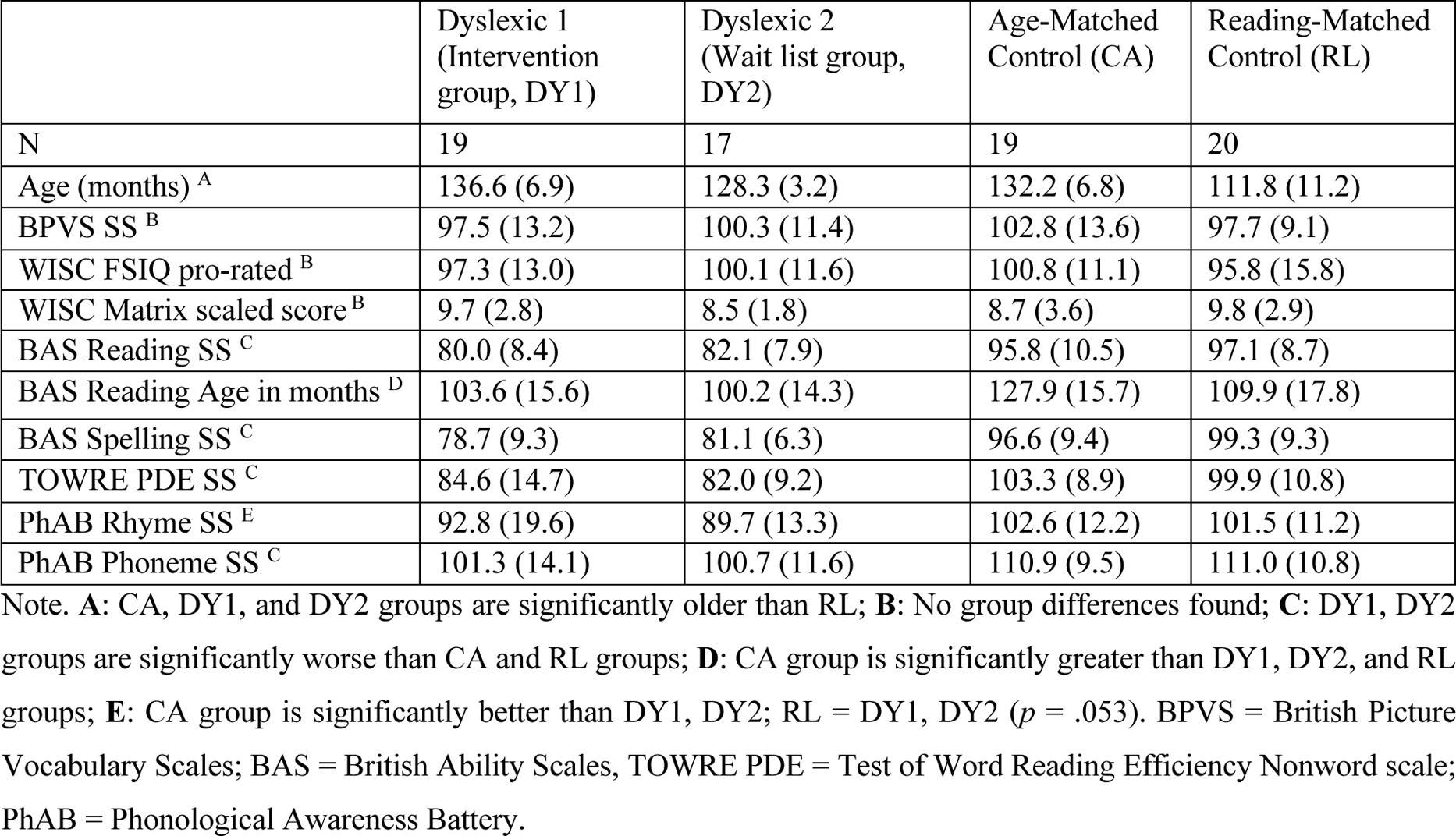
Details of the participating children.

### 2.2. Stimuli

The auditory stimuli were 20 multi-syllabic words or phrases listed in Table 2. The computerized speech stimuli were created from a native female speaker of standard Southern British English, and sampled at a 44.1-kHz rate with 24-bit resolution. Stimuli were presented at the same sampling rate of 44.1 kHz, and were chosen to ensure that they included a mixture of stressed and unstressed syllables in different positions within the words or phrases.

**Table 2.** Words and phrases used in the study.

|  |  |  |  |  |
| --- | --- | --- | --- | --- |
| Aladdin | Bambi | Bob The Builder | Dr Who | Flushed Away |
| Harry Potter | Horrid Henry | Madagascar | Over The Hedge | Scooby Doo |
| Shaun The Sheep | The Cat In The Hat | The Gingerbread Man | The Jungle Book | The Lion King |
| The Muppets | The Simpsons | The Smurfs | Wallace And Gromit | Winnie The Pooh |

### 2.3. Experimental set-up

Participants were seated beside the experimenter in front of a laptop in a quiet room. They were presented with the 20 auditory stimuli through a MATLAB-based task. To ensure that the participant understood the task correctly, the experimenter and participant carried out 2 practice trials before the main experiment using their names. For example, they might say “Now we are going to play a game about listening to rhythm patterns in words. Word rhythm goes with the syllables. Listen to my name: fi-O-na / AN-ge-la / ANN-a-bel. Can you hear the rhythm?” The experimenter then repeated their name and the child’s name, tapping on the table with a finger. “Okay, now I’ve got some new ones for you to listen to on the computer”.

During presentation of each experimental stimulus, the AE of the stimulus and a picture providing a memory prompt were presented on the task layout (see Figure 1). The children were instructed to listen to the stimulus and to repeat three times what they heard, and they were encouraged to try to match their response line to the visual AE display on each occasion. The visual AE matching was intended to provide some feedback concerning their performance. Their responses were recorded by a microphone, with an expected 60 responses per child. Each response was recorded over a time interval of 3 seconds. After the stimulus was presented, the target picture (Figure 1A) and the stimulus AE (Figure 1B) remained on the main layout. As the child repeated a target stimulus, the AE of the child’s latest response appeared on the main layout overlaying the AE of the target stimulus (Figure 1C).

### 2.4. Speech features

Two multi-syllabic speech features, the AE and the pitch contour, were investigated. Before applying the similarity analyses, the quiet parts at the beginning and at the end of the auditory responses were removed manually. The AE was calculated as the absolute value of Hilbert transform, using function *hilbert* (.) from MATLAB, of the speech signal, and then low-passed filtered at 50 Hz. Pitch contour was also calculated, using function *pitch* (.) from MATLAB, as the estimates of the fundamental frequency over time window of 50 ms for the speech signal. Both speech features were resampled for children’s utterances to have the same length as the corresponding target stimuli.

### 2.5. Similarity measures

Two objective measures were used to quantify the similarity between the child’s response AE and the child’s response pitch contour and the AE and pitch contour of the target stimuli. They were the Pearson correlation (*r*) and Mutual Information (*MI*). The correlation between variables *X* = {*x*_1_, *x*_2_, …, *x*_3_} and *Y* = {*y*_1_, *y*_2_, …, *y*_3_} is calculated as:

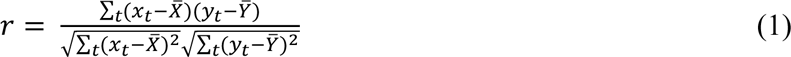

where 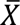 and 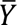 are average for variable *X* and *Y*, respectively. *MI* between two random variables is defined as a measure of information that one random variable gives about the other one, and is calculated as (Cover, 1999):

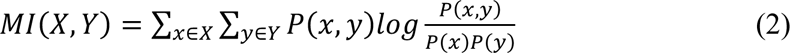

where *P*(*x*) and *P*(*y*) are the marginal distributions of variables *X* and *Y*, respectively, and *P*(*x*, *y*) is the joint distribution of these variables. Here marginal and joint distributions were estimated using the Gaussian kernel estimator:

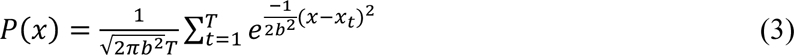

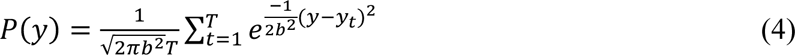

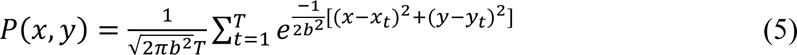

where *b* refers to the bandwidth parameter that tunes the kernel function (Qiu, Gentles, & Plevritis, 2009).

### 2.6. Statistical analysis

As children were encouraged to produce three responses to each target, the mean scores (separately for *r* and *MI*) were computed for the 1st, 2nd, and 3rd responses respectively for each participant. This does not mean that all children had three responses for each single trial. For example, one child might have only two responses for the first trial (’Aladdin’) and three responses for the rest of the trials (19 trials). This child would still have three averaged repetition scores, as the average score for the third repetition was computed by averaging over 19 trials. Separate analyses were carried out for the AE and pitch contour metrics respectively, since the derived measures of similarity (*r* and *MI*) were computed differently mathematically and were also computed using different acoustic parameters (AE versus pitch contour).

If the opportunity to repeat the target items three times led to a learning effect across responses, then the *r* and *MI* values would be expected to increase parametrically from the 1st to the 3rd response for each metric. In such a case, a main effect of repetition could be expected. *A priori*, we expected the children with dyslexia to be poorer at producing the AEs of the target phrases. In this case, a main effect of group could be expected for the AE*-r and* AE*-MI* analyses. This was explored statistically using 4 repeated measures ANOVAs, with group as the between-subjects factor and repetition as the repeated measure (repetition 1, repetition 2, repetition 3). The dependent variable in each case was each similarity metric (AE*-r,* AE*-MI,* pitch contour*-r,* and pitch contour*-MI)*. Prior to analysis, the data were first explored for normality using the Explore function in SPSS v27, to check they did not violate criteria for parametric analyses. Two outliers were identified as values more than a 1.5 interquartile range below the lower quartile of the population data, a DY1 child who had a very small AE-*r* value on their second repetition, and a CA child who had a very small pitch contour-*r* value on their third repetition. These two data points were excluded before applying repeated measures ANOVAs to the dataset.

## 3. Results

Figure 2 shows the speech production data by group for the *AE* and Figure 3 shows the speech production data by group for pitch contour. As noted, *a priori*, we expected the children with dyslexia to show worse speech production in terms of similarity to the target pronunciations for the AE metrics (*r* and *MI*). We did not make an *a priori* prediction for the pitch contour metrics. It was also thought possible that repeated practice and the visual feedback of viewing the AE could improve repetition accuracy from repetition 1 to repetition 3 for the AE measures. The repeated measures ANOVAs supported the first expectation regarding group differences, but not the second expectation regarding potential AE learning effects during the experiment. The repeated measures ANOVAs showed a significant effect of repetition for all *four* analyses, AE-*r*, *F*(2,140) = 14.8, *p* < .001; AE-*MI*, *F*(2,142) = 9.3, *p* < .001; pitch contour-*r*, *F*(2,140) = 11.4, *p* < .001; pitch contour-*MI*, *F*(2,142) = 18.4, *p* < .001. Post-hoc Newman-Keuls tests showed that for each metric, the children showed significantly better accuracy for repetition 2 compared to repetition 1, but significantly less accurate repetition of the targets when comparing repetition 3 to repetition 2. Accordingly, there is no strong evidence for learning effects during the experiment, as improved similarity regarding target production did not increase parametrically across repetitions. Rather, all groups performed best on their second repetition. The data also suggest that the visual feedback of seeing the AE did not have a systematic effect on performance, as repetition effects were similar for both the AE and the pitch contour analyses. As will be recalled, pitch contours were not depicted visually during the task. It is difficult to explain why children got significantly worse on their third repetition, however it could have been fatigue with the task.

**Fig 2.**
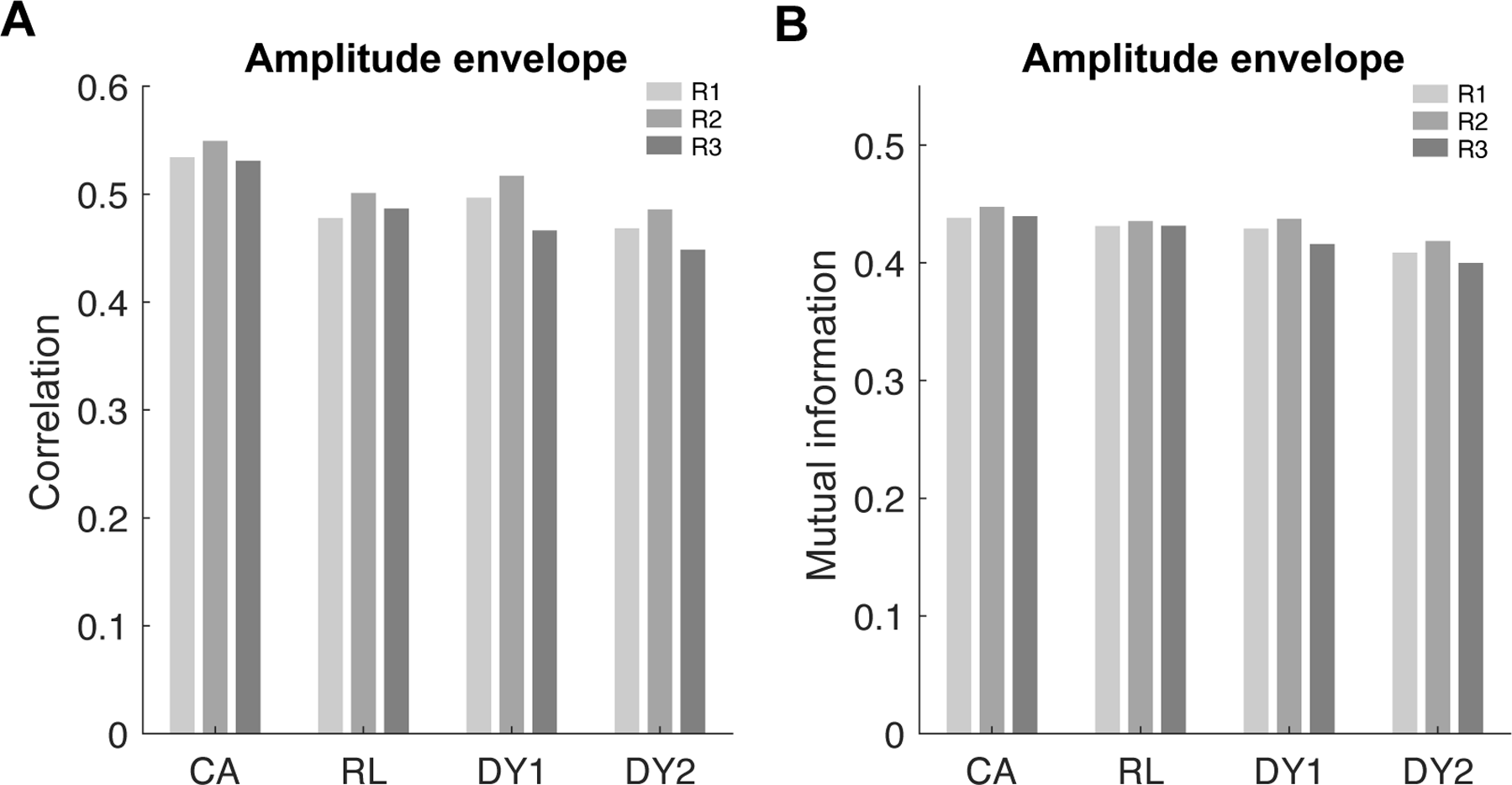
Performance for the four groups (CA, RL, DY1: children with dyslexia receiving intervention, DY2: children with dyslexia waiting for intervention) in amplitude envelope matching by repetition (R1, R2, R3) for (A) the correlation between stimuli envelopes and the envelopes of corresponding responses, and (B) the mutual information between stimuli envelopes and the envelopes of corresponding responses.

**Fig 3.**
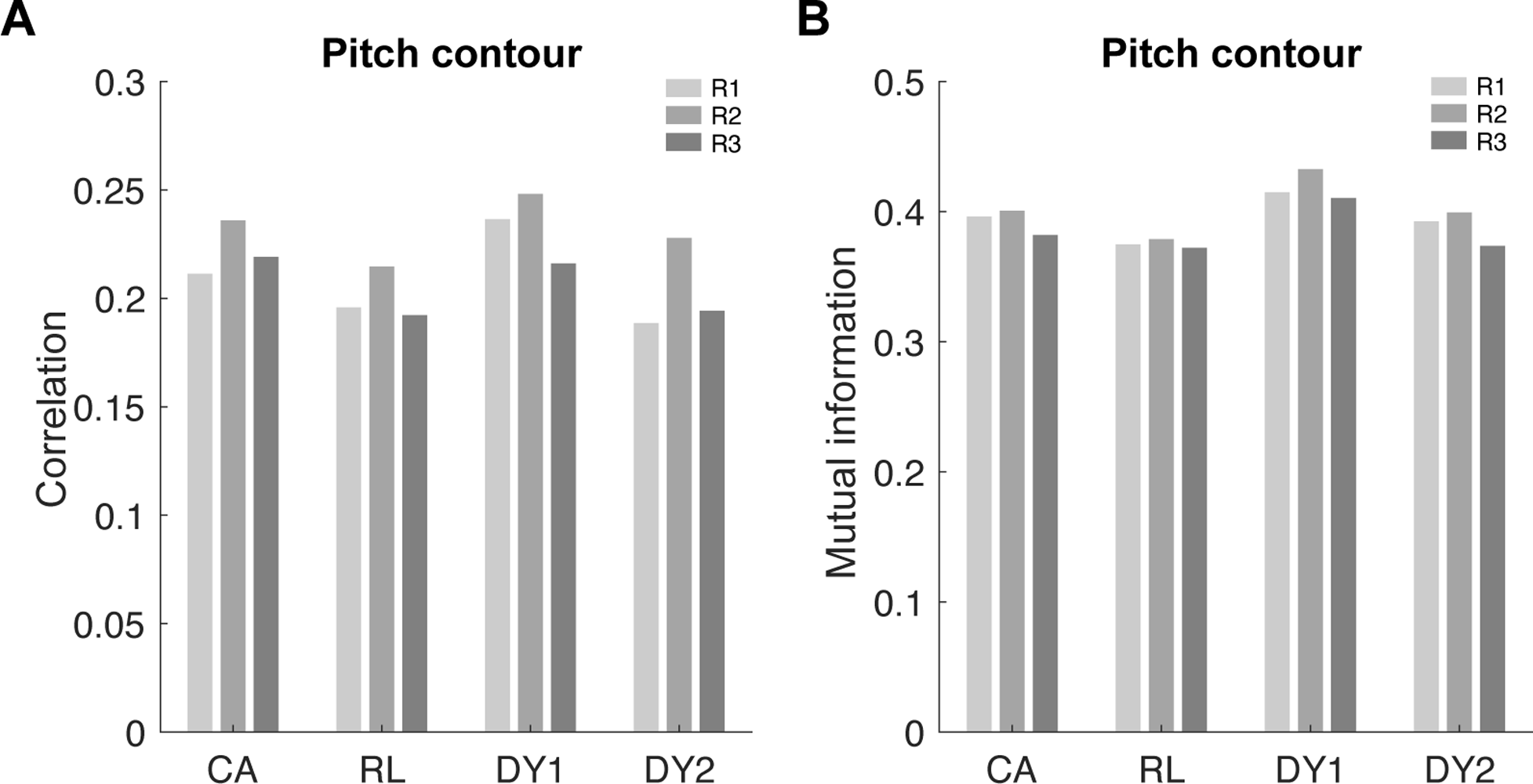
Performance for the four groups (CA, RL, DY1: children with dyslexia receiving intervention, DY2: children with dyslexia waiting for intervention) in pitch contour matching by repetition (R1, R2, R3) for (A) the correlation between stimuli pitch contours and the pitch contours of corresponding responses, and (B) the mutual information between stimuli pitch contours and the pitch contours of corresponding responses.

A priori, we had expected the AE analyses to show significant effects of group. However, three of the four repeated measures ANOVAs showed a significant main effect of group, AE-*r*, *F*(3,70) = 4.7, *p* = .005; AE-*MI*, *F*(3,71) = 3.4, *p* = .022; pitch contour-*MI*, *F*(3,71) = 3.3, *p* = .026. The exception was pitch contour-*r*, *F*(3,70) = 1.7, *p* = .25. None of the repeated measures ANOVAs showed a significant interaction between repetition and group, all *p*’s > .05. Exploring the group effects for the AE metric first, which were predicted *a priori*, post-hoc tests (Newman-Keuls, one-tailed) for the AE-*r* similarity metric showed that both groups of children with dyslexia were significantly worse than the CA children at producing AEs that matched the targets, DY1 versus the CA children, *p* = .038, DY2 children (those waiting for an oral intervention) versus the CA children, *p* = .001. The DY1 children were also significantly better than the DY2 children at producing matching AEs, *p* = .039. In addition, the RL group were producing less similar AEs to the targets than the older CA group as estimated by the *r* similarity metric, *p* = .005, as would be expected.

For the AE-*MI* metric, the post-hoc tests showed that the DY2 children were again producing significantly less similar AEs to the target pronunciations compared to the CA group (*p* = .002), and were also producing significantly less similar AEs compared to the younger RL group (*p* = .013). The DY1 children did not differ significantly from either control group for the *MI* metric, but were again significantly better at matching the target pronunciations than the DY2 group, *p* = .041. Accordingly, children with dyslexia (DY2 children) who were not yet receiving an oral intervention designed to improve their phonological awareness of multi-syllabic words and phrases were significantly worse at producing an accurate AE for the target phrases than younger RL-matched children for the *MI* similarity metric, suggestive of a fundamental difficulty. Children with dyslexia (DY1 children) who were receiving an oral intervention designed to improve their phonological awareness of multi-syllabic words and phrases did not show impaired AE performance compared to typically-developing control children for the *MI* metric.

Regarding potential group differences in pitch contour, there were no significant group differences for the pitch contour-*r* metric, indicating no difference in pitch contour similarity to the target between groups. For the *MI* metric, post-hoc tests (Newman-Keuls, two-tailed) showed that the DY1 children were significantly *better* at pitch matching than the younger RL control children, *p* = .003, and showed a trend to also be better than the CA control children, *p* = .075. The DY1 children were also significantly better than the DY2 children, *p* = .044. The DY2 children did not differ from either the CA or RL control children. These data indicate that producing accurate pitch contours for multi-syllabic words and phrases is not impaired in children with dyslexia. They also suggest that the oral intervention being received by the DY1 children may have been affecting their pitch contour production.

Figure 4 depicts both the AEs and the spectrograms for the stimulus “Aladdin”, along with the corresponding responses from participants from each group who were chosen to illustrate the overall effects depicted in Figure 2 (CA, RL, DY1, DY2). For comparison, pitch contours from the same child participants are also depicted.

**Fig 4.**
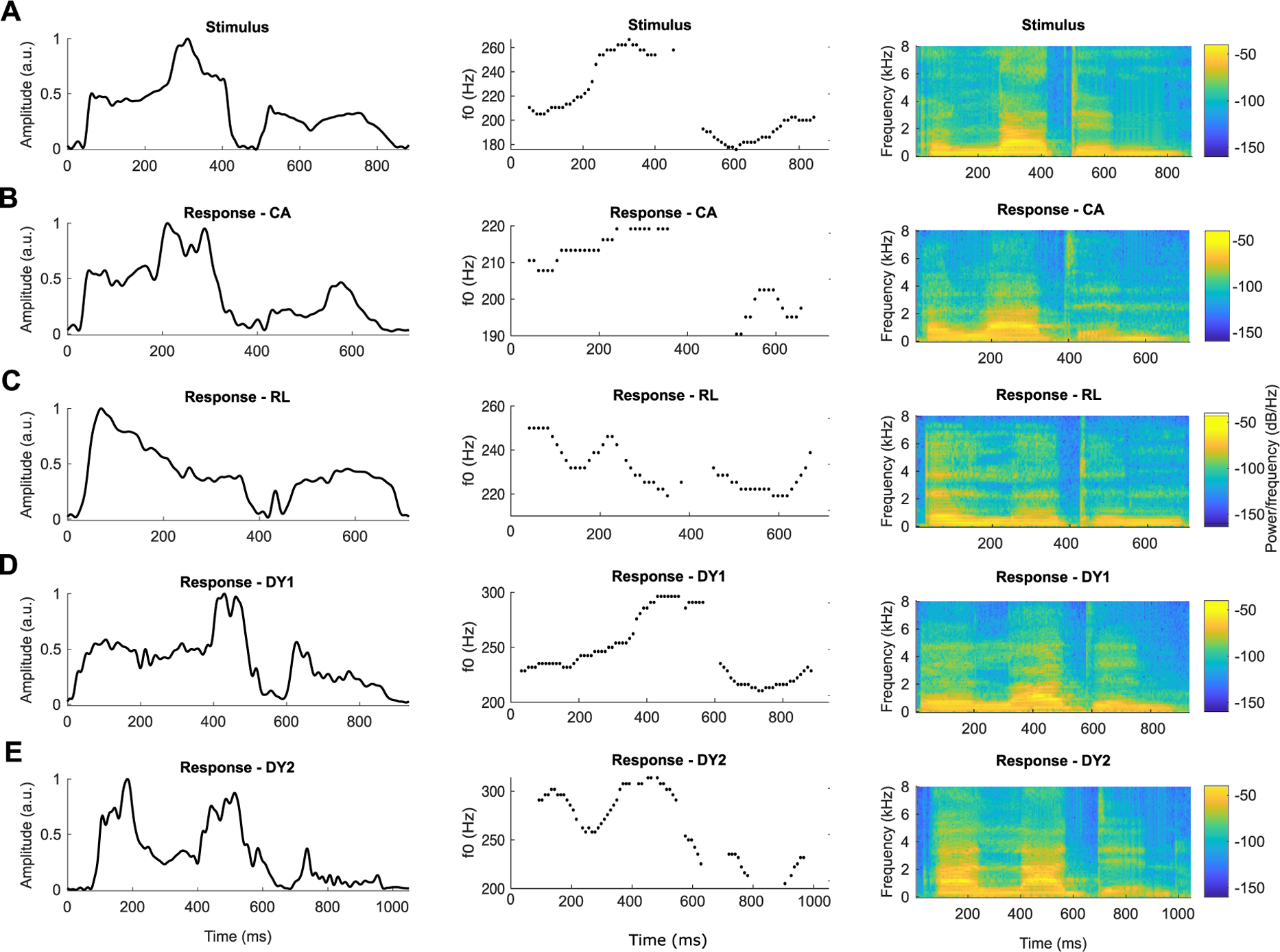
Examples of the amplitude envelopes (left), pitch contours (middle) and spectrograms (right) produced by selected individual children in each group for the target stimulus “Aladdin”. Panels in row A depict the target spoken by an adult, panels in row B are for an example CA child, panels in row C are for an example RL child, panels in row D are for an example child with dyslexia receiving intervention (DY1), and panels in row E are for an example DY2 child (a child with dyslexia waiting for intervention). Note that for the correlation and mutual information analyses, the child’s speech productions were resampled to have the same length as the corresponding target stimuli. Also note that the pitch estimates are depicted only for the analysis windows with the harmonic component.

## 4. Discussion

Children with dyslexia are known to show impaired perception of the syllable stress patterns in multisyllabic words and phrases (e.g., Goswami et al., 2010, 2013). Here we measured whether children with dyslexia also show impaired production of syllable stress patterns in multisyllabic words and phrases. On the basis of TS theory, we predicted impaired production of AE information, but not of pitch contour. Using a simple and novel speech copying task, we indeed found robust difficulties in the accurate production of the AEs of familiar multi-syllabic words by children with developmental dyslexia. Children with dyslexia who were not yet receiving an oral rhythmic intervention designed to support their phonological awareness of multi-syllabic words (DY2 group) were particularly impaired regarding AE similarity to the targets, as they also performed significantly more poorly than younger RL-matched control children for the *MI* similarity measure. These children (DY2 group) also performed significantly more poorly in matching the AE than the children with dyslexia who were receiving the oral intervention designed to support their phonological awareness of stress patterns (DY1 group). This suggests that the 18-session oral intervention within which the stress copying task was embedded for the DY1 group does help children with dyslexia to perceive and produce speech rhythm patterns. Nevertheless, the intervention appeared to have modest effects regarding AE copying, as the DY1 children were still significantly poorer at producing AEs that matched the targets than the CA children using the *r* similarity measure. Regarding the AEs that were produced by children with dyslexia, the depictions of speech output for individual children provided in Figure 4 suggest that children with dyslexia were not producing an envelope of amplitude (intensity or loudness) differences between strong (stressed) and weak (unstressed) syllables that were similar in overall AE shape to the oral targets.

Of particular note, all groups performed at similar levels in terms of matching the pitch contours of the target items (see Figure 3). This rules out any specific motoric or articulatory difficulties that could explain the impaired AE performance shown by the children with dyslexia. Indeed, inspection of the individual example data depicted in Figure 4 suggests that the pitch contours produced by the children with dyslexia (DY1 and DY2 groups) were highly similar to the target pitch contours, with the DY1 group producing significantly more similar pitch contours to the targets than the younger RL-matched controls. Pitch contours capture the overall changes in fundamental frequency (f0) as speakers alternate between stressed and unstressed syllables and is a core aspect of intonation (Ladd, 1980). The relative success of children with dyslexia in producing pitch contours may help to explain why the speech production difficulties of children with dyslexia regarding the AE appear to pass unnoticed by their listeners and teachers.

Accurate pitch contour production in dyslexia would not be consistent with a prior auditory theory of developmental dyslexia, in which difficulties relating to rapid spectro-temporal changes that reflect phonemes rather than the relatively slow spectro-temporal changes reflected in the AE are proposed as the central auditory deficit (Tallal, 2004). Interestingly, school-aged children with dyslexia have been thought to have difficulties in nonword repetition tasks (Melby-Lervåg & Lervåg, 2012, for a meta-analysis), where performance is typically scored in terms of phonemes correct. To our knowledge, no published studies of nonword repetition score children’s performance in terms of the accurate reproduction of stress patterns. If nonword repetition skills were scored at the level of syllable stress patterns, the nonword repetition literature in dyslexia may be more consistent than reported in the meta-analysis by Melby-Lervåg & Lervåg (2012). Melby-Lervåg & Lervåg’s meta-analysis demonstrated that the nonword repetition difficulties at the phoneme level found in the reviewed studies of school-aged children actually depended on the proportion of children included who had developmental language disorder rather than dyslexia. Consistent with TS theory and a rhythm-based analysis, nonword repetition skills in pre-school (hence pre-reading) samples are known to be related to general rhythmic skills, such as tapping to a beat (Kalashnikova et al., 2021). Further, rhythm copying skills at age 5 predict non-word repetition at age 7 (Moritz et al., 2013).

The demonstration of impairments in word production at the level of syllable stress and the AE instead supports an alternative auditory theory of dyslexia, TS theory (Goswami, 2011). As noted earlier, TS theory is based on perceptual difficulties with both speech and non-speech rhythm related to sensory impairments in perceiving amplitude rise time and amplitude modulation. These rhythmic difficulties can be indexed behaviourally by tapping tasks, musical rhythm discrimination tasks and syllable stress perception tasks (e.g., Wolff, 2002; Thomson & Goswami, 2008; Huss et al., 2011; Goswami et al., 2013). As it is based on rhythm, TS theory is the only theory of dyslexia that would predict impaired speech production of the stress patterning in multi-syllabic words and phrases at the level of the AE. The demonstration that such impairments are indeed present in children with dyslexia suggests that *both* speech input and speech output processes in dyslexia are impaired with respect to AE information. The data are also convergent with the pioneering longitudinal toddler work with at-risk samples (Lambrecht Smith et al., 2010; Smith et al., 2008). Here, differences in speech production at the syllable level were measurable long before a diagnosis of developmental dyslexia could be obtained.

There are many null results in the dyslexia speech production literature, but most such studies have explored the *phonetic* level of speech representation. Here negative results (no differences between groups) are interpreted as showing no difficulties in speech production. For example, when researchers used a range of tasks designed to measure ‘phonological grammar’ (the typically unconscious and phonetic processes that are applied in speech production, such as voicing assimilation), French dyslexic adults performed as well as control adults (Szenkovits, Darma, Darcy, & Ramus, 2016). In voicing assimilation in French, the voicing feature may spread backwards from obstruents or fricatives to the preceding consonant, so that for instance “cape grise” [ka**p**griz] becomes [ka**b**griz] (grey cloak). Szenkovits et al. (2016) reported that French adults with dyslexia produced exactly the same voicing assimilations as other adults. Similarly, French adults with dyslexia were able to produce non-native phonemes (Korean plosives) as accurately as non-dyslexic adults (Soroli, Szenkovits, & Ramus, 2010). These data were interpreted as being inconsistent with the phonological theory of dyslexia (Ramus & Szenkovits, 2008). The current data suggest instead that such studies have been analysing speech production measures at the wrong linguistic level. Speech production measures that tap into speech rhythm and the accurate production of syllable stress patterns may be more sensitive to individual differences between participants with and without dyslexia. The finding that toddler speech production differences were measurable at the syllable level rather than the phoneme level also suggests that linguistic analyses testing the core phonological deficit model of dyslexia require analyses at the supra-segmental level (Smith et al., 2008).

The current child data have some limitations. It is unfortunate that group matching of the CA and DYS groups was affected by the Pandemic, and future studies should compare groups of dyslexic and control children of the same age. The participating children were also relatively old when they received the speech copying task. Given the data from at-risk toddlers, it would also be important to compare groups of younger children. A further limitation is that the visual feedback of displaying the AE did not seem to affect performance for any group in a systematic way. Although repetition effects were significant, this arose in part because all children got worse on their third repetition, and furthermore similar repetition effects were found for the pitch contour analyses even though pitch contour was not depicted on screen. Accordingly, different forms of feedback than a visual depiction may be more useful in helping children with the copying task.

The data also have novel practical and clinical implications. First, it may be possible to identify risk for dyslexia early in development by adding multi-syllabic measures of speech production to current diagnostic tests. Although children show considerable individual variability in speech production measures, the addition of such measures could enable diagnosis and intervention long before children enter school and begin to experience literacy failure. Secondly, the data support the efficacy of dyslexia interventions that work at the level of syllable stress and speech rhythm (Bhide, Power, & Goswami, 2013; Cancer et al., 2020; Flaugnacco et al., 2015). It is likely that developmental effects will be optimized the younger that such interventions can occur (Kalashnikova, Burnham, & Goswami, 2021). Thirdly, acoustic interventions that address the sensory problems in perceiving AE structure in dyslexia are likely to be beneficial. For example, amplifying AE rise time cues in natural speech may help the dyslexic brain with speech processing and consequently facilitate development of a well-specified phonological system (Van Hirtum, Ghesquiere, & Wouters, 2021; Van Hirtum, Moncada-Torres, Ghesquiere, & Wouters, 2019). Finally, it is notable that language acquisition is thought to begin with speech rhythm, with stressed syllables providing clues to word onset in languages like English (Cutler & Carter, 1987; Mehler et al., 1988). The demonstration that speech production in dyslexia is impaired at the level of syllable stress patterns suggests that the dyslexic experience of oral language is different from that of typically-developing children from infancy onwards. This could suggest that early interventions to ameliorate dyslexia should enhance the acoustic rhythm patterns that are naturally exaggerated in infant-directed speech (Leong, Kalashnikova, Burnham, & Goswami, 2017). Continuing to use infant-directed speech for longer in early life with at-risk infants may help to decrease the difficulties in perceiving and producing syllable stress patterns that characterize children with dyslexia.

## Ethics statement

All children and their parents gave informed consent for the study in accordance with the Declaration of Helsinki, and the study was approved by the Psychology Research Ethics Committee of the University of Cambridge.

## Acknowledgments

The authors would like to thank all the children, families and schools involved in the study. This research was funded by a grant awarded to UG by the Fondation Botnar (project number: 6064) and a donation to UG from the Yidan Prize Foundation. The sponsors played no role in the study design, data interpretation nor writing of the report. GDL was supported by Science Foundation Ireland under the Career Development Award 15/CDA/3316 (EL) and under Grant Agreement No. 13/RC/2106_P2 at the ADAPT SFI Research Centre at Trinity College Dublin. ADAPT, the SFI Research Centre for AI-Driven Digital Content Technology, is funded by Science Foundation Ireland through the SFI Research Centres Programme.

## Author contributions

M.K. contributed to conceptualization, methodology, data analyses, visualization, and writing – original draft; G.D.L. contributed to conceptualization, methodology, and writing – review and editing; F.G. contributed to investigation and writing – review and editing; A.W. contributed to investigation and writing – review and editing; A.M. contributed to investigation; U.G contributed to funding Acquisition, project administration, supervision, conceptualization, methodology, and writing – original draft.

## Data availability statement

The analyses were performed using MATLAB and SPSS. Any data (including costume codes) with the exception of the children’s original voice recordings can be shared upon request.

